# Expression reflects population structure

**DOI:** 10.1101/364448

**Authors:** Brielin C Brown, Nicolas L. Bray, Lior Pachter

## Abstract

Population structure in genotype data has been extensively studied, and is revealed by looking at the principal components of the genotype matrix. However, no similar analysis of population structure in gene expression data has been conducted, in part because a naïve principal components analysis of the gene expression matrix does not cluster by population. We identify a linear projection that reveals population structure in gene expression data. Our approach relies on the coupling of the principal components of genotype to the principal components of gene expression via canonical correlation analysis. Futhermore, we analyze the variance of each gene within the projection matrix to determine which genes significantly influence the projection. We identify thousands of significant genes, and show that a number of the top genes have been implicated in diseases that disproportionately impact African Americans.

**Author Summary:** High dimensional, multi-modal genomics datasets are becoming increasingly common, which warrants investigation into analysis techniques that can reveal structure in the data without over-fitting. Here, we show that the coupling of principal component analysis to canonical correlation analysis offers an efficient approach to exploratory analysis of this kind of data. We apply this method to the GEUVADIS dataset of genotype and gene expression values of European and Yoruban individuals, finding as-of-yet unstudied population structure in the gene expression values. Moreover, many of the top genes identified by our method have been previously implicated in diseases that disproportionately impact African Americans.

## Introduction

Genes mirror geography to the extent that in homogeneous populations, individuals can be localized to within hundreds of kilometers purely on the basis of their genotype [1,2,3]. Population structure in genotypes is revealed via projection of single nucleotide polymorphism (SNP) data onto the first few principal components of the population-genotype matrix. The principal components space, which is a lower-dimensional distinguished subspace of the high-dimensional data, is computed by a procedure called principal components analysis (PCA). While PCA has been successful in revealing population structure from SNP data, it does not identify such structure in some other genomic data types. For example, in the case of gene expression data, PCA has not revealed obvious population signatures (Figure 1A, [4]). Here we show that although the first two principal components of expression data do not capture population structure, there are other projections that do. One approach to finding such projections is the coupling of dimension reduction to correlation maximization after adjustment for confounding. This approach, utilizing PCA and canonical correlation analysis (CCA), has been used to effectively analyze the relationship between gene expression and copy number variation [5]. The method is implementable via singular value decomposition and is therefore also efficient. We apply it to finding population structure in expression data, thereby further highlighting the combination of PCA and CCA as a powerful approach to integrative analysis of genomics data.

**Figure 1:**
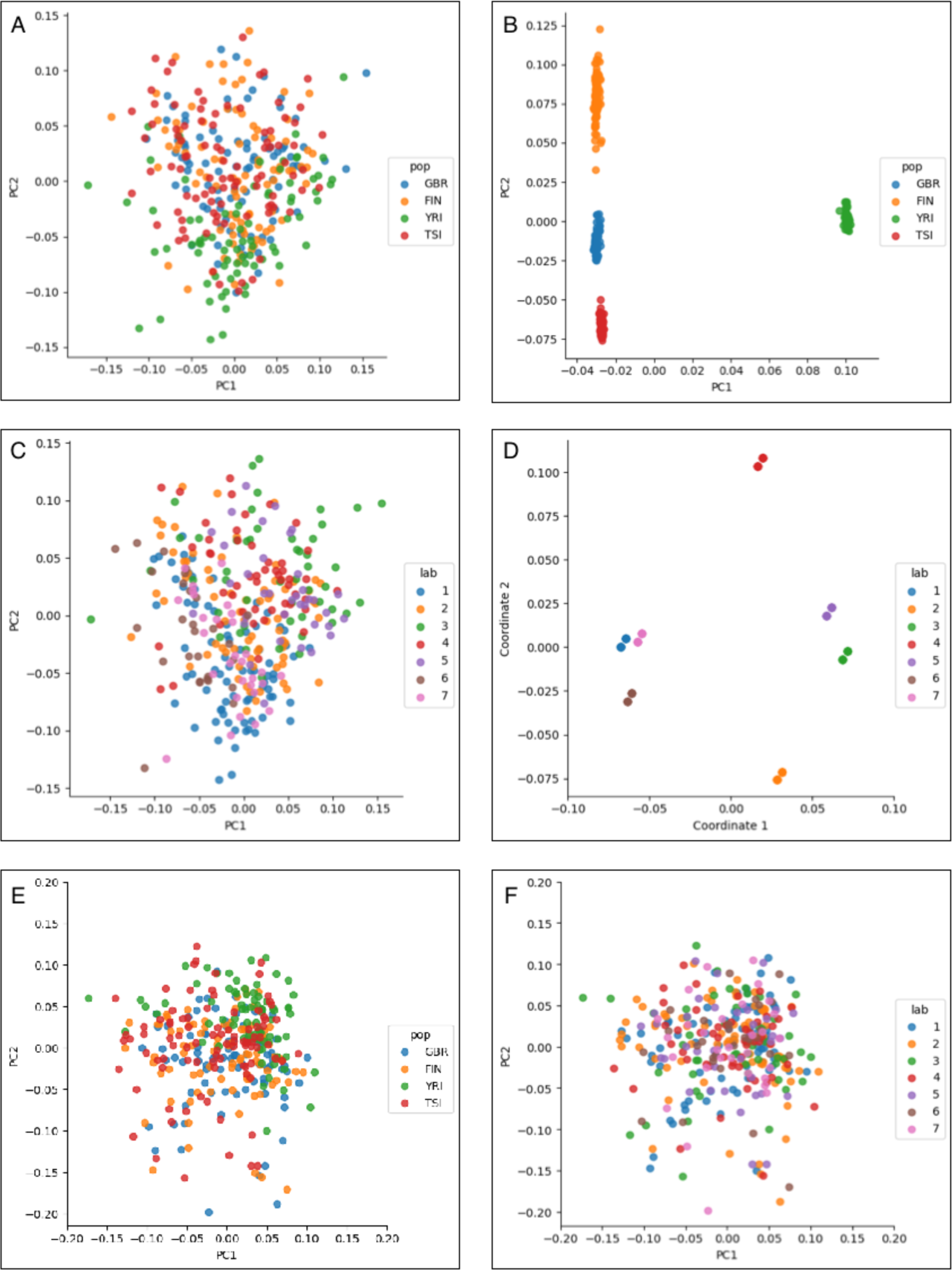
(A) PCA of the expression matrix fails to reveal clustering by population, whereas (B) PCA of the genotype matrix reveals clear clustering by population. (C) Coloring of samples by batch reveals that PC1 and PC2 are being partly defined by batch source. (D) Projection into the space learned by a regression model of gene expression from batch and gender identity revealas strong clustering in the data. (E) PCA of the batch-corrected expression matrix. (F) The corrected data no longer cluster by batch.

As an optimization procedure, PCA can be viewed as the projection of data onto the lower-dimension subspace that minimizes the average distance of the data to its projection. This is algebraically equivalent to finding the lower-dimensional subspace that maximizes the variance of the projected data. This statistical view of PCA helps to explain why PCA of expression data might not reveal population structure: even if such structure is present in the data, it may not lie on the directions of maximal variance. CCA is a widely used method for joint analysis of heterogeneous data and provides a linear-algebraic mechanism for identifying shared structure among a pair of datasets. Given a pair of data matrices, CCA finds maximally correlated linear combinations of the columns of each matrix [6]. We show that CCA applied to the PCA projections of expression and genotype data identifies a projection of the expression data that reveals population structure, which we validate via a leave-one-out cross validation experiment.

To validate our method, we examined population structure in expression data from the Genetic European Variation in Health and Disease (GEUVADIS) project [7], which consists of RNA-seq data obtained from lymphoblastoid cell lines derived from whole-genome sequenced individuals belonging to five distinct populations. The GEUVADIS data has been extensively studied [7], yet our analysis reveals structure not previously examined in this well-characterized dataset.

## Results

A naïve PCA analysis of the GEUVADIS expression data (Figure 1A) shows that unlike genotype data (Figure 1B), there is no clear clustering of individuals by population. This result is consistent with other analyses of expression data, in which population structure is not detected by PCA [4]. To understand the sources of variation that could explain the first and second principal component axes, we labeled the individuals according to the lab where they were sequenced (Figure 1C). This provides some insight into the sources of variation. For example, samples from Lab 3 are distinctly separated from Lab 1. We therefore proceeded to correct for confounding by regressing the gene expression matrix on a matrix of potentially confounding variables and taking the residual (Figure 1D—F, see Methods). We note that it is also possible to correct for confounding using CCA by exploiting the relationship between CCA with categorical data and linear discriminant analysis [8] (Supplementary Methods, Supplementary Figure 1).

Next, we examined whether coupling of expression data to genotype data could identify a projection that reveals population structure. To do this we performed PCA followed by CCA on the batch-corrected expression matrix and the genotype matrix (see Methods). The resulting CCA projection of expression data (Figure 2A), reveals distinct population patterns in the data, although not as clearly as the resulting PCA of the genotype data (Figure 1B). We validate this observation by performing a leave-one-out cross-validation experiment, where we remove each individual from the dataset and show that the reconstruction error of the model on the held out point is close to the error in the training set, and that the principal components of the reconstructed gene expression matrix show similar population patterns (Figure 2B). Moreover, the CCA projection is indexed by linear combinations of genes, which can be understood to discriminate individuals based on expression signatures. That is, genes with high variance in the CCA expression projection (see Supplemental Methods) have expression distributions that segregate based on patterns in the genotype PCs, which we interpret to represent population structure [1,2,3]. After correction for correlated multiple testing using the Benjamini–Hochberg–Yekutieli procedure [9], we identified 3,581 genes with significant scores at FDR 5%, indicating that population structure within gene expression data is pervasive.

**Figure 2:**
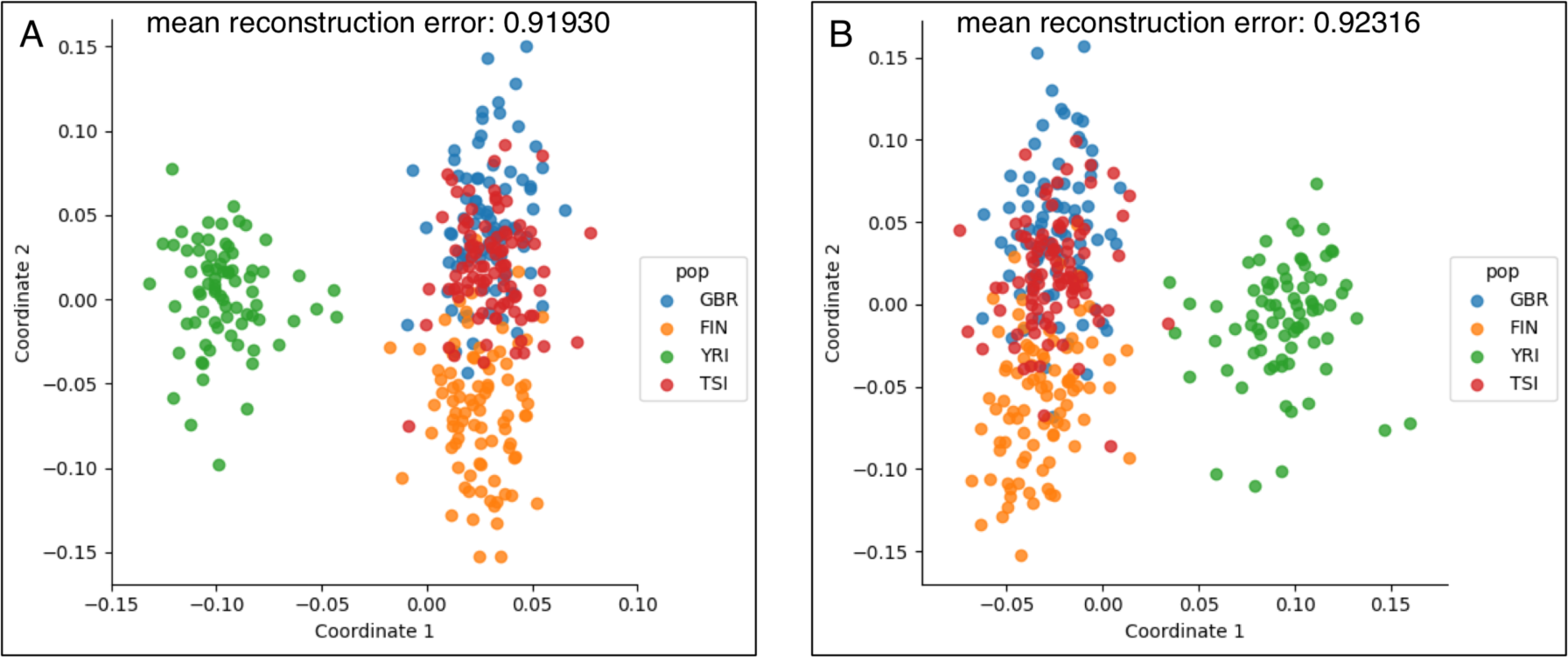
(A) A CCA projection of the batch-corrected expression matrix that shows that expression reflects population structure. While the individuals, labeled according to their population, do not cluster as clearly as with genotype data (Figure 1B), there is clear population structure in the CCA projection of the batch-corrected expression data. (B) A leave-one-out crossvalidation experiment showing that individuals are approximately projected to their populations of origin even when the projection matrix is learned without their expression or genotype data. The mean re-construction errors in (A) the left-in samples and (B) the held-out samples are similar and overlayed on top of the Figure. The first two canonical correlations are 0.963 and 0.766.

The three genes with largest z-score in this analysis were LATS-2, EIF4EBP2 and STX7 (Figure 3). All three genes display increased expression in the YRI population, highlighting the possibility of relating population expression differences directly to disease phenotypes such as discussed in [10], where EIF4B is shown to be associated with vascular disease in African Americans. In addition, LATS-2 is a known tumor suppressor gene, and it is well known that African Americans have significantly worse outcomes at every stage of cancer treatment [11]. Other notable genes identified with this method include PSPH, NEDD4-2 and ARHGEF11 (all with p<1e-07). PSPH was examined in [12] where it was found to be the gene with the highest degree of differential allelic expression. The eQTL rs6700, which is associated with expression of that gene, is an ancestry informative marker [13]. NEDD4-2, which is associated with hypertension [14], contains SNPs such as rs945508 which are similarly associated with ancestry. Allelic variants in ARHGEF11 are associated with kidney disease in mouse models [15], which has higher prevalence in the African-American community. While we view the identification of such genes as important, we caution that African-Americans also experience substantial structural inequality in healthcare, which confounds this analysis [16]. We also note that while genes such as NEDD4-2 must also have substantial genetic/epigenetic regulation that is linked to population differences, the projection-associated genes identified by our method does not produce that information. Indeed, its power to detect genes associated with population structure comes by virtue of requiring only one test per gene and being agnostic to the source of regulation. While a complete analysis of population-associated expression differences is beyond the scope of this paper, these examples suggest that our method should be a powerful approach for directly identifying genes whose expression associates with population.

**Figure 3:**
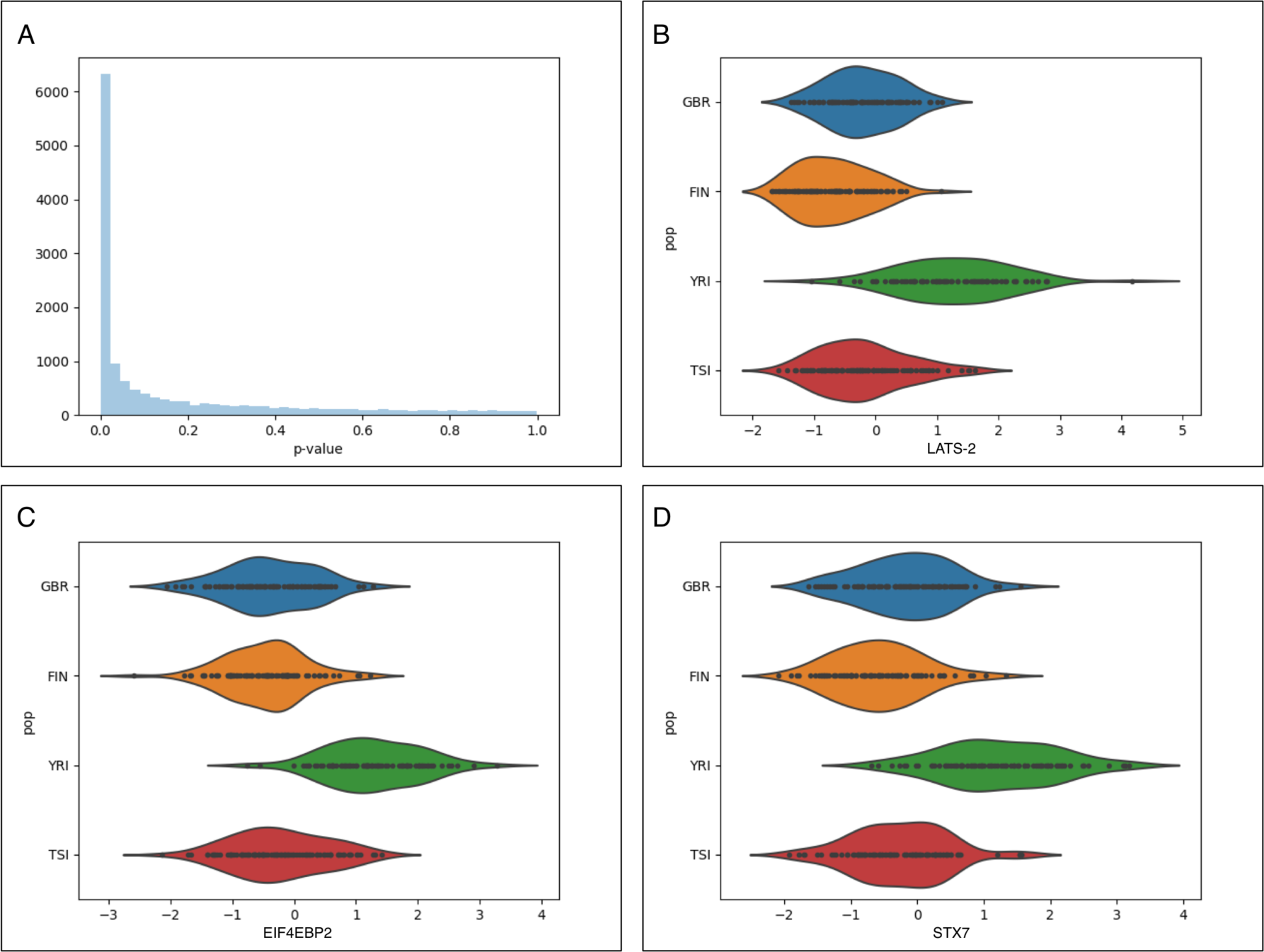
(A) The p-value distribution for tests that the variance of each gene in the projection is greater than the null shows a large number genes with significant scores in the PCA+CCA projection. The expression distributions by population for the three genes with highest z-scores are shown in (B) the LATS-2 gene, (C) the EIF4EBP2 gene, (D) the STX7 gene.

## Conclusion

The identification of population structure in expression data suggests that it should be interesting to extend population genetic methods such as [17] to population transcriptomics. The example of joint analysis of expression and genotype data can be extended to include other datatypes via an extension of CCA to more than two matrices [8,18,19,20], and the coupling of PCA to CCA could also be extended to a hierarchical factor analysis method. Importantly, the coupling of PCA and CCA is not the only projection that reveals population structure. For example, connecting the principal components using linear regression gives similar visualizations (Supplementary Methods, Supplementary Figure 2). The choice of model should reflect the variance structure of the data, which here we have deliberately remained agnostic to. Ultimately, it is important to identify the optimal model for inference.

While we believe the extensions describe above will be interesting to pursue, our analysis and that in [5] shows that PCA+CCA is a useful and rapid approach to exploratory analysis of heterogeneous data. As the generation of large-scale, high-dimensional, multi-modal genomics datasets becomes more commonplace [20, 21, 22], we expect the combination of PCA and CCA to become as common as PCA is today.

## Methods

We obtained genotype data of the Phase 1 1000 genomes individuals in PLINK format [23] from cog-genomics [See Data and Software Availability]. GEUVADIS project RNA-seq reads were downloaded from the European Nucleotide Archive (accession number ENA: ERP001942). In the analyses performed we omitted the CEU population because it has been previously found to display biased expression patterns due to the age of the cell line [24]. Importantly, this bias affects every CEU sample and therefore cannot be corrected for traditional methods of handling confounding.

There are 343 individuals with genotype data from 1000 genomes phase 1 and corresponding RNA-seq data from GEUVADIS in the FIN, GBR, TSI and YRI populations. We quantified the transcript abundances of these individuals using kallisto [25] with the GENCODE v27 protein coding transcript sequences and annotation. The GENCODE v27 annotation contains 95,659 transcripts. We omitted all transcripts with mean transcripts per million (TPM) less than 0.1 across the quantified samples, leaving 58,012 transcripts. We then used the GENCODE v27 annotation to obtain gene level quantifications by summing transcript quantifications in TPM units. Finally, we removed genes in the MHC region and on non-autosomal chromosomes. This left 14,070 genes for analysis. The Phase 1 1000 genomes genotypes contain 39,728,178 variants. We filtered indels, variants with minor allele frequency (MAF) less than 5%, and non-biallelic SNPs leaving 6,785,201 SNPs for analysis. Finally, we quantile-normalized the expression matrix, and centered and scaled each gene quantification vector to have mean 0 and variance 1. In the following analyses, we chose to keep 27 principal components of expression and 11 principal components of genotype, while analyzing the first two canonical components. However we note that our results are stable under different choices of numbers of components (Supplementary Figure 3, Supplementary Table 1).

To remove batch effects from the expression matrix, we one-hot encoded the lab identification vector, and then added a column for sample gender [26], resulting in a 343 x 7 matrix of potentially confounding variables. We then regressed each gene expression vector on the confounding matrix and used the residual expression vector for all further analysis. Next, we computed principal components of the genotype matrix using PLINK and principal components of the corrected expression matrix using the eigendecomposition of the sample correlation matrix. Finally, we computed the canonical variables between the top principal components of the genotype and corrected expression matrices (see the Supplementary Methods for details on the linear algebra).

To verify that we did not over-fit in estimating coefficients using CCA, we performed leave-one-out cross validation. We removed each of the 343 individuals one-by-one from the dataset, re-calculated the principal components of the genotype and expression matrices, and reestimated the canonical variables and bases. We then projected each held out individual into the resulting CCA gene expression subspace. After this process, for each individual, we plotted the first two principal components of the re-constructed expression matrix to verify the individual clusters by population (see the Supplementary Methods for details of the how the projection was performed). Furthermore, we calculated the in-sample and out-of-sample reconstruction error as the squared Frobenius norm of the original and reconstructed data points, and verified that it was similar for both left-in and held-out samples.

Finally, we asked which genes had significant variance for the CCA gene expression projection. We computed the variance of each gene in the projection, and calculated significance via a permutation test with 10 million permutations. In each iteration, we shuffled the genotype principal components and recomputed the variance explained. The p-value derived from this test is the number of times the permuted score is greater than the observed score, divided by the number of permutations (see the Supplementary Methods for details of how the variance is computed). We further estimated a z-score for each gene as the difference between the estimated and mean permutation variance divided by the variance of the permuted variance.

## Data and software availability

The software used to produce the analyses is on GitHub. We provide a package of tools for computing the projections and estimating gene significance, as well as a Snakemake file [27] that can be used to completely reproduce the analysis, from data acquisition to figure generation.

- Analysis software: https://github.com/pachterlab/PCACCA/
- Gencode v27 transcripts: ftp://ftp.sanger.ac.uk/pub/gencode/Gencode_human/release_27/gencode.v27.pc_transcripts.fa.gz
- Gencode v27 GTF: ftp://ftp.sanger.ac.uk/pub/gencode/Gencode_human/release_27/gencode.v27.annotation.gtf.gz
- GEUVADIS RNA-seq reads: ftp://ftp.sra.ebi.ac.uk/vol1/ERA169/ERA169774/fastq
- 1000 genomes genotypes: https://www.dropbox.com/s/k9ptc4kep9hmvz5/1kg_phase1_all.tar.gz

## Acknowledgements

The authors would like to thank Shannon McCurdy for invaluable feedback on this manuscript.

